# Topography of distance-modulated multisensory object location encoding in mouse area RL

**DOI:** 10.64898/2026.02.04.701688

**Authors:** Yue Zhang, Uwe Lewin, Alessandro La Chioma, Tobias Bonhoeffer, Mark Hübener

**Affiliations:** Max Planck Institute for Biological Intelligence, Martinsried, Germany; Graduate School of Systemic Neurosciences, Ludwig-Maximilians-Universität München, Martinsried, Germany

## Abstract

The spatial arrangement of an animal’s sensory environment is encoded by spatial codes of the external world, often simultaneously in different modalities. How these modalities are brought together in a universal spatial map and how potential differences are reconciled during multisensory integration remains unclear. To tackle this question, we systematically mapped object location receptive fields (OLRFs) across different sensory modalities in area RL, a pivot of visuo-tactile integration within mouse posterior parietal cortex, using a custom high-resolution three-dimensional multisensory stimulation system.

Most RL neurons have three-dimensional receptive fields, with many exhibiting OLRFs across visual, tactile (i.e. vibrissal), and bimodal stimulus conditions. Comparing OLRFs across modalities revealed that distance strongly shapes visuo-tactile integration, reflected in the shift of bimodal OLRFs from tactile-dominant to visual-dominant with increasing distance. Distance also affects the linearity and spatial mode of integration. At near distances, neurons integrate object location information sublinearly from overlapping regions of visual and tactile OLRFs. At far distances, information is integrated supralinearly from regions covered by OLRFs of either modality. This distance dependency extends to cortical organization, leading to a distance tuning gradient along which the spatial, modality, and integration properties of RL neurons are aligned. Together, our findings suggest distance as a key factor in visuo-tactile integration, reflecting that the limited reach of whiskers makes visual information more important for farther objects. In parallel, the system shifts from a localization dominated mode for nearby objects to a detection dominated mode for farther objects.

## Introduction

Deeply different, yet all senses dwell within one shared external space. To interact coherently with the external world, the brain must integrate object location information distributed across sensory modalities.

Such integration is inherently complex because object location is encoded in fundamentally different ways across modalities. First, each sensory modality samples distinct regions of external space. For instance, rodent whiskers densely sample peri-facial but not distant space, whereas vision assesses distant space but is poor at localizing close-by objects (such as those obscured by the nose or other body parts).

Second, the reference frames of object location differ across modalities and vary across processing stages: Visual representation of space begins with retinotopic maps, where each neuron encodes a location on the two-dimensional retinal surface, with information on the third dimension of space, depth, added at subsequent stages^1–7^. In contrast, tactile representation of space in the rodent whisker system starts with somatotopic maps, where object location is initially tied to whisker identity, and later generalized into a head-centered spatial representation^8–11^. To form an integrated representation of space, the brain must reconcile differences in spatial coverage and reference frames. Understanding multimodal integration therefore requires identifying the spatial reference frames used by each modality, and the mechanisms resolving the modality-specific differences.

The posterior parietal cortex (PPC) has emerged as a key neural substrate for multimodal integration in mammals^12^. Within mouse PPC, area RL, located between primary visual (V1) and barrel cortex (S1), serves as a hub for visuo-tactile integration^13^. As a multisensory region with extensive reciprocal connections to V1 and S1, RL contains unimodal neurons, which respond exclusively to either visual or tactile stimuli, and bimodal neurons that respond to both^13–17^. The bimodal neurons integrate visual and tactile input in both supralinear and sublinear ways, suggesting a complex mechanism for multisensory integration^13,18^. Together, these findings highlight RL’s pivotal role in visuo-tactile integration.

A recent study showed that many neurons in RL are tuned to binocular disparity, with a clear preference for very close objects, which likely fall within whisker reach^2^. The coherent visual and tactile representation of upper and lower space in RL described recently further implies the emergence of a unified egocentric framework in the multimodal coding for near space^16^. However, this co-alignment could alternatively arise from the aligned arrangement of two-dimensional somatotopic and retinotopic maps, without necessarily requiring a representation of three-dimensional near space. Resolving this ambiguity requires systematic mapping of object location tuning across modalities.

In this study, we addressed this question using a high-resolution stimulation system for precise mapping of receptive fields under unimodal visual or tactile as well as bimodal conditions, in behaving mice. Our results reveal a primarily co-aligned egocentric reference framework across modalities, also reflected in cortical topographical maps. We show that visuo-tactile integration is profoundly distance-dependent, affecting not only modality preference, but also the mechanism and linearity of integration.

## Results

To investigate how object location information in mouse near space is extracted and integrated from visual and tactile inputs in area RL, we developed a motorized multimodal stimulus system to map object location receptive fields (OLRFs) in RL neurons (Figure 1A). This system delivers small objects accessible to both vision and whiskers to precise 2D and 3D locations, enabling controlled, repeatable presentation under different sensory conditions. We achieved independent presentation of tactile and visual cues by placing awake, head-fixed mice in darkness to remove other visual inputs, and using a motorized whisker mask to block tactile access (Figure 1B, 1C). This approach enables separate measurement of modality-specific OLRFs and direct comparison of receptive field properties across modalities. A vertical bar served as the visual and tactile stimulus for 2D mapping, while a small sphere enabled mapping of object location preference along all axes in 3D.

**Figure 1.**
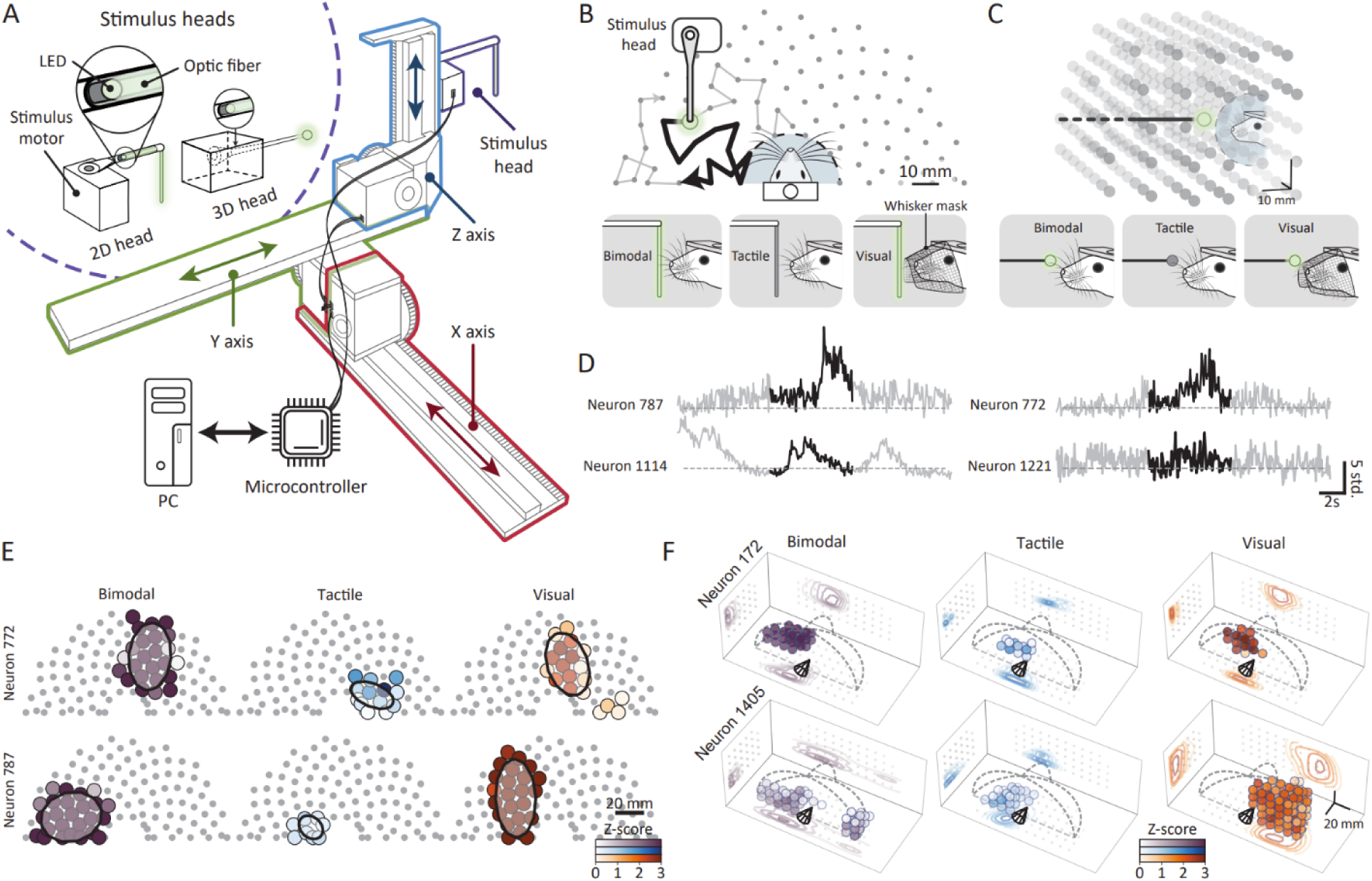
Multimodal stimulus system for mapping object location receptive fields (OLRFs) in mouse area RL. **A.** Schematic of the custom-built visuo-tactile stimulus system comprising three components: (1) a 3-axis motorized translation stage, (2) interchangeable stimulus heads for 2D and 3D stimulation, and (3) on-board microcontroller interfacing with PC running custom stimulation software. **B-C.** Stimulus paradigms for 2D (**B**) and 3D (**C**) object location mapping. Gray dots indicate stimulus positions sampled during each recording session. Blue shaded region denotes collision protection zone (15 mm radius). Bottom panels illustrate three experimental conditions: *Bimodal* - stimulus is illuminated and accessible to whiskers; *Tactile* - stimulus is dark, but remains accessible to whiskers; *Visual* - stimulus is illuminated, but whiskers are masked to prevent tactile contact. **D.** Z-scored calcium activity traces from four example neurons during 2D stimulus presentation. Gray and black segments highlight neural activity during bimodal stimulation moving along the corresponding trajectory shown in panel **B**. **E.** 2D object location receptive fields (OLRFs) from two example neurons shown in **D** across different stimulus conditions. Large colored dots indicate statistically significant OLRF components (see Methods and Figure S1), colored by the corresponding mean calcium activity (Z-scores) at each stimulus position. Receptive fields are fitted with ellipses for quantification. Small gray dots mark all tested stimulus positions. **F.** 3D OLRF examples from two neurons across stimulus conditions. Scatter points indicate significant OLRF components colored by mean calcium activity. Dashed curves outline the total stimulation volume. Contour plots show firing rate distributions estimated from neural responses to stimulus positions projected onto XY/XZ/YZ planes (gray dots) using Gaussian kernel density estimation.

We designed a continuous stimulus protocol that allows efficient measurement of OLRFs for all neurons recorded across all modality combinations with dense spatial sampling (∼10 mm) in less than 14 minutes for 2D stimuli and less than 18 minutes for 3D stimuli (see Methods for details). To avoid spatiotemporal correlations due to continuous stimulation, the protocol iterates through a fixed set of near space locations in a semi-random, trial-unique order determined by a heuristic traveling salesman algorithm (Figure 1B, 1C). The OLRFs are computed by reverse-correlating calcium activity acquired by two-photon imaging with stimulus trajectories across all trials (five trials for 2D stimuli; three trials for 3D stimuli; examples in Figure 1D). Significant OLRF components were detected with a two-step approach (see Methods and Figure S1).

Using this approach, we successfully measured 2D OLRFs of 3,782 cells (more precisely, regions of interest, or ROIs; see also Methods) in area RL and the RL/V1 border region from seven mice, and 3D OLRFs of 2,351 cells from five mice. In total, 86.7% of cells for 2D stimulation and 81.2% for 3D stimulation showed significant OLRFs under at least one condition. The majority of the discovered OLRFs were spatially localized and spanned all measured dimensions (Figure 1E-F), suggesting that most neurons are indeed tuned to object locations relative to an external reference frame. In other words, these neurons encode object positions in space rather than the retinotopic or somatotopic position or other lower-dimensional stimulus features correlated with location such as size or luminance.

Differences in the geometry of sensory epithelia and modality-specific processing might lead to variations in how object location is encoded across modalities. To understand how visual and tactile spatial codes differ, and to what extent these differences are resolved during integration, we compared OLRF properties across all stimulus conditions (Figure 2A, 2K). Some neurons responded exclusively to object location in a single modality, while others integrated both visual and tactile inputs (Figure 2A, 2K).

**Figure 2.**
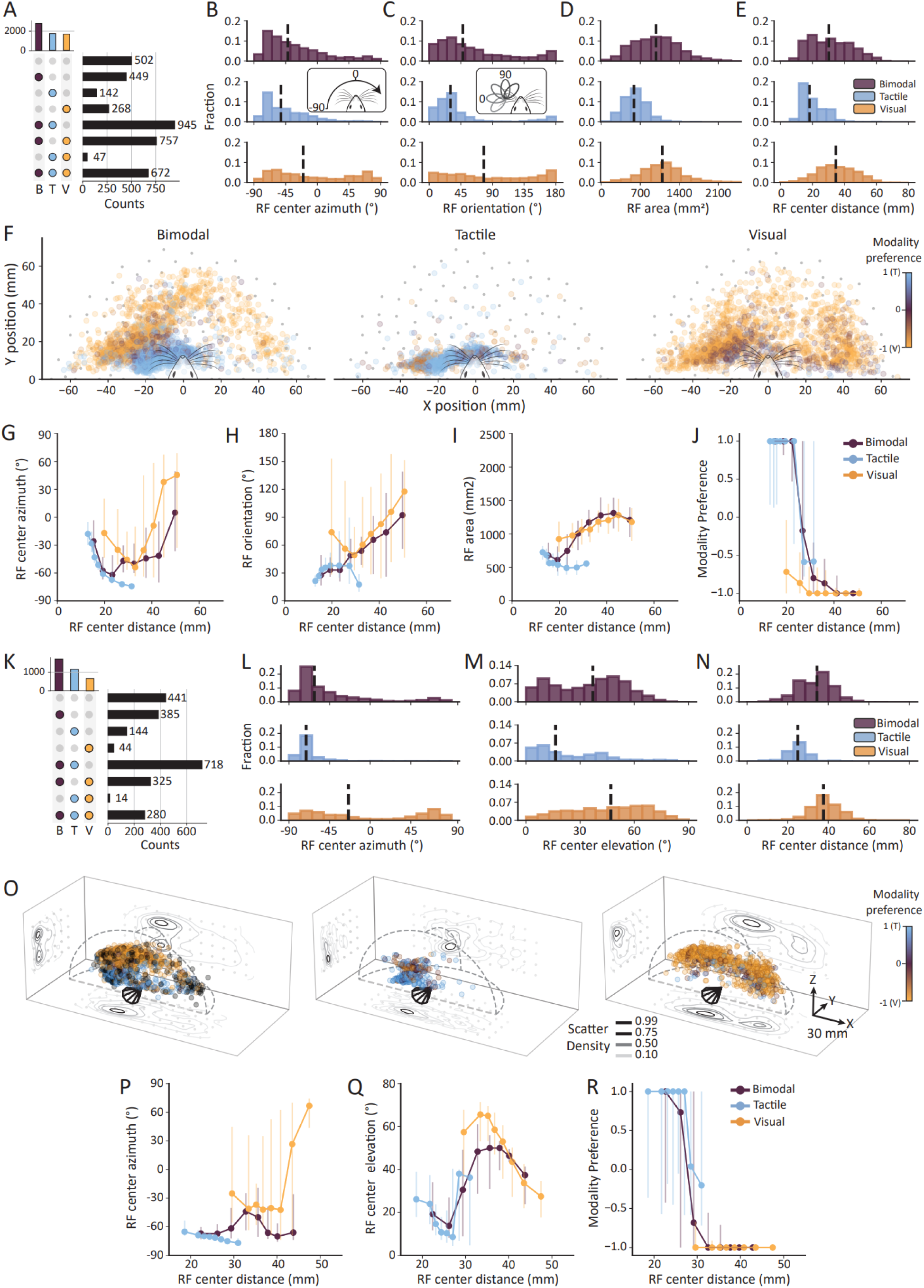
OLRF properties of RL neurons vary systematically with preferred object distance. **A.** UpSet plot quantifying neurons with significant 2D OLRFs across three stimulus conditions (B: bimodal, T: tactile, V: visual). Top bar chart shows total OLRF counts per condition. Colored circles in bottom chart: neurons with significant OLRFs under the corresponding conditions; neuron counts for each group are indicated by horizontal bars. **B-E.** Histograms of 2D OLRF center azimuth (**B**), receptive field orientation (**C**), covered area (**D**), and center distance (**E**) under different stimulus conditions (purple: bimodal; blue: tactile; yellow: visual). The vertical dashed lines indicate medians of distributions. The distributions of all OLRFs properties differ significantly across stimulus conditions (Mann–Whitney U test; p<0.001). **F.** Spatial distribution of 2D OLRF centers on the egocentric near-space plane where the X/Y axes correspond to left-right/anterior-posterior directions. The color of the large scatter points represents the neuron’s modality preference (see Methods). Small gray dots indicate all stimulus positions. **G-J.** 2D OLRF center distances versus OLRF center azimuth (**G**), receptive field orientation (**H**) and covered area (**I**) on the egocentric plane, and modality preference (**J**). The nodes and error bars correspond to median and the 25^th^ −75^th^ percentiles. **K.** UpSet plot for neurons with significant 3D OLRFs across stimulus conditions, following the same conventions as in panel **A**. **L-N.** Histograms of 3D OLRF center azimuth (**L**), elevation (**M**), and distance (**N**) under different stimulus conditions. **O.** Scatter plots of 3D OLRF center distribution in mouse near space. The color of the scatter points represents neurons’ modality preference. Cone-shaped polygons indicate mouse head position and orientation. Contour lines represent scatter density estimated with Gaussian kernels (normalized to [0,1], 1 indicates the peak density), projected onto cardinal planes (gray dots: stimulus position projections). **P-R.** The preferred distances of 3D OLRFs versus azimuth (**P**), elevation (**Q**), and modality preference (**R**). The nodes and error bars in **P-R** represent median and the 25^th^ −75^th^ percentiles.

To quantitatively compare the spatial extent and orientation of OLRFs across modalities, we fitted ellipses (2D) or ellipsoids (3D) respectively to each OLRF. Histograms in Figure 2B-E show that OLRFs for different modalities have distinct properties (Mann-Whitney U test; p < 0.001 for all condition pairs in each panel). Tactile OLRFs were smaller, more oriented along the left-right axis, and clustered in the contralateral proximal region of the mouse near space. Visual OLRFs were larger and more evenly distributed across the near space, while bimodal OLRFs showed intermediate properties (Figure 2B-E). These trends were consistent for both 2D and 3D OLRFs (Figure 2L-N), indicating systematic differences in object location encoding across modalities.

Systematic differences in population-level OLRF statistics across stimulus conditions could result from modality-specific receptive fields in individual neurons, as illustrated by example neurons (Figure 1E, 1F). Alternatively, they could arise from correlations between OLRF properties and neurons’ modality preferences, resulting in the recruitment of different neuron populations with distinct OLRFs across conditions. To distinguish between these possibilities, we calculated a modality preference index based on each neuron’s mean responses to tactile and visual stimuli within its OLRFs (see Methods). The distributions of modality preference for 2D and 3D OLRFs (Figure 2F, 2O) revealed a clear distance dependence: neurons with OLRFs closer to the animal were more tactile-dominant, while those with more distal OLRFs were more visual-dominant. This distance effect is most pronounced in the bimodal stimulus condition, but is also present in visual and tactile conditions (Figure 2J, 2R). The consistency across conditions suggests that the distance-modality relationship results from intrinsic integration properties of RL neurons, rather than simply from the sensory inputs available.

Other receptive field properties also varied with distance in a modality-dependent manner (Figure 2G-I, 2P-Q). In all cases, bimodal OLRFs at near distances resembled tactile OLRFs, while those at farther distances resembled visual OLRFs. The transition from tactile-like to visual-like OLRFs typically occurred at a distance of 25–30 mm in both 2D and 3D conditions, which is approximately the maximum whiskers reach^14^. These findings indicate that distance-dependent encoding of object location is a general property of RL neurons involved in visuo-tactile integration.

The primary visual and somatosensory cortices are organized as two-dimensional retinotopic or somatotopic maps^19,20^. Our data show how RL neurons encode object locations in a three-dimensional reference frame. This raises the questions of whether and how distance, the missing dimension in the somatotopic and retinotopic maps, is integrated into the topographic organization of RL. To assess this across animals, we established a shared anatomical framework in two steps: First, we aligned cortical regions from different recording sessions within each animal using blood vessel landmarks (Figure 3A-B). These aligned regions were then registered across animals using azimuth maps derived from bimodal 2D OLRFs (Figure 3C, see Methods for details). RL-V1 boundaries were identified from retinotopic maps of individual animals obtained with intrinsic optical imaging as references.

**Figure 3.**
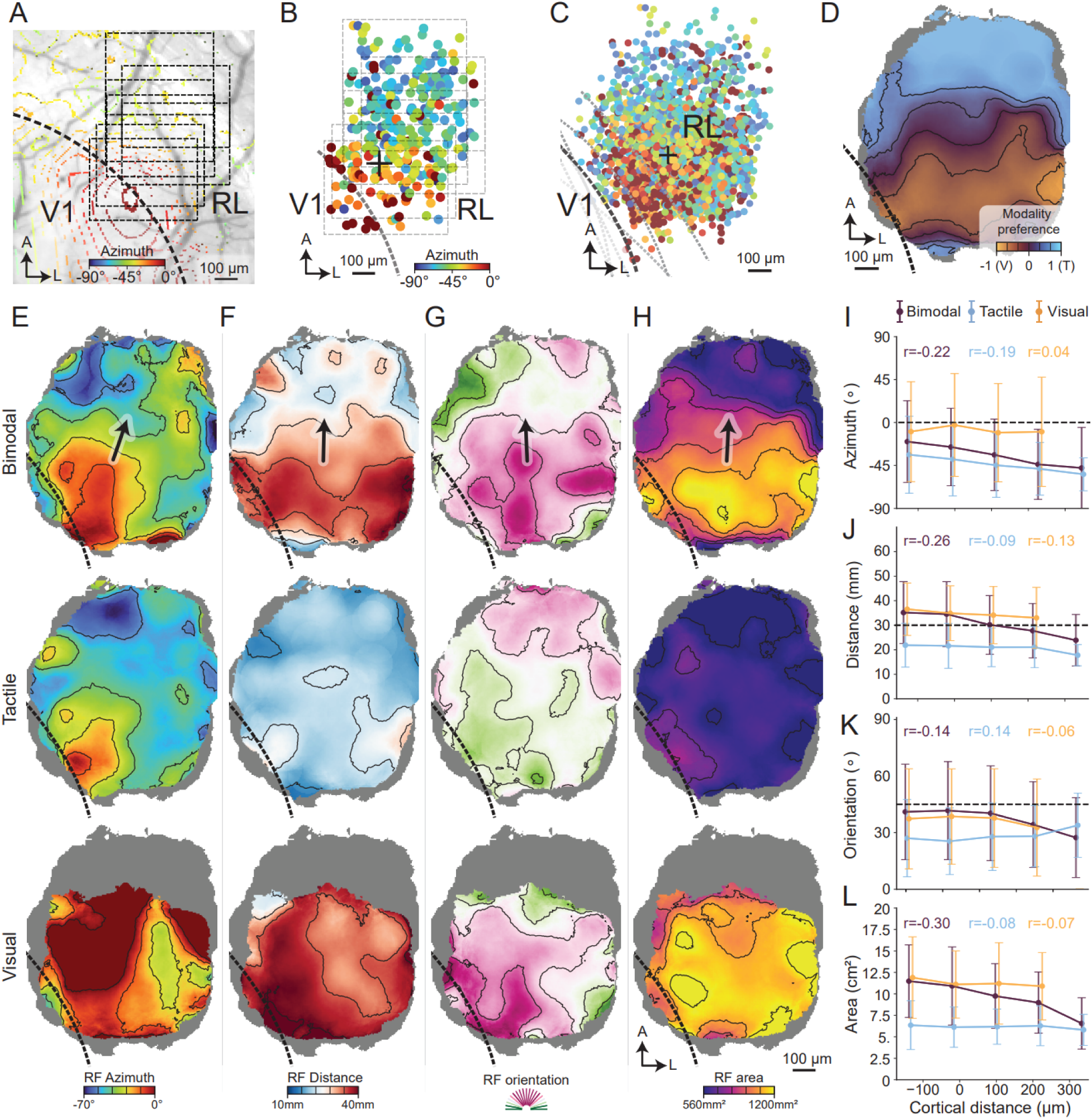
Cortical topography of 2D OLRFs reveals organized maps of visual and tactile spatial preferences. **A.** Overlay of retinotopic azimuth map and cortical surface vasculature from intrinsic optical signal imaging in an example mouse. Dashed squares: the region recorded during multiple two-photon calcium imaging sessions. Dashed line: V1-RL boundary. **B.** The OLRF azimuth map obtained from the two-photon calcium imaging sessions (dashed squares) shown in **A**. The “+” marks the median position of neurons tuned to medial visual field (azimuth > −15°). Colored dots represent the anatomical locations of neurons with significant bimodal OLRFs, colored by azimuth preference of OLRF centers. **C.** Alignment of azimuth maps across animals by offsetting individual maps with median positions of near vertical median tuned neurons (the “+” mark; see Methods for detailed alignment procedure). Gray dashed lines: individual mouse V1-RL boundaries based on intrinsic optical signal imaging retinotopic maps; Black dashed lines in C-H indicate the estimated average RL-V1 boundary across animals. **D.** Smoothed modality preference map showing the spatial distribution of visual-dominant, tactile-dominant, and balanced multisensory neurons across RL. Gray regions indicate insufficient data (n < 10 cells within the 180 μm neighboring region). Contour lines show iso-preference boundaries at intervals marked on the color bar. **E-H.** Smoothed feature maps of OLRF azimuth (**E**), distance (**F**), orientation (**G**) and area (**H**) under bimodal (top), tactile (middle), and visual (bottom) stimulus conditions. Black arrows indicate the principal gradient direction vector in the corresponding bimodal map of each column. Gray regions indicate insufficient data (n < 10 cells within the 180 μm neighboring region). Contour lines indicate iso-value boundaries at intervals marked on the corresponding color bars. **I-L.** OLRF azimuth (**I**), distance (**J**), orientation (**K**), and area (**L**) changes along the principal gradient direction vectors in **E-H** (x axis; 0 is the projection of the origins in **E-H)**. Points and error bars indicate the mean ± standard deviation. Distance bins with less than 100 cells were excluded from the plots. The Pearson r for each condition is labeled in color, as indicated in the legend in **I**. A, anterior; L, lateral.

Within this framework, we characterized the cortical organization of 2D OLRF properties across bimodal, visual, and tactile conditions. Spatial patterns were visualized as Gaussian-smoothed feature maps, with pixel colors representing distance-weighted averages of nearby neurons’ OLRF properties (Figure 3D-H; also see Methods).

The resulting maps revealed systematic gradients across multiple OLRF properties. Bimodal OLRF azimuth in RL showed a continuous gradient, shifting from medial to peripheral positions when moving away from V1 (Figure 3E, top), consistent with the retinotopic organization. We also observed a gradient of modality preference shifting from visual to tactile dominance over the same span (Figure 3D), consistent with previous studies^13,16^. Similar gradients were observed for bimodal OLRF distance and orientation (Figure 3F-H). Here, orientation refers to how the receptive field is arranged in physical space around the animal, rather than how it is oriented on the retina. The OLRFs of neurons close to V1 tended to be located farther in the mouse near space, larger, and oriented caudal-rostrally, while those farther from V1 were more often found in the proximal regions of mouse near space, smaller and oriented along the medial-lateral axis.

Maps based on visual or tactile OLRFs often diverged from bimodal maps in regions dominated by the other modality. As a result, bimodal maps resembled visual maps near V1 but aligned more closely with tactile maps far from V1 (Figure 3I-L). This shift produced steeper spatial gradients across bimodal maps compared to unimodal maps. Similar trends were also observed in 3D OLRF maps (Figure S2). It is worth noting that the lack of an azimuth gradient in the visual maps in Figure 3E and S2C is largely attributable to the emergence of a substantial number of ipsilateral OLRFs under visual-only conditions (see also Figure 2F and 2O). These findings demonstrate that the topographic organization of RL neurons is shaped by multimodal integration, rather than simply mirroring the properties of single-modality sensory inputs.

The partial misalignment between bimodal and unimodal maps suggests that visuo-tactile integration of object information involves more than a simple combination of unimodal inputs. This raises the question of how this integration is implemented in individual neurons, especially in those with spatially offset visual and tactile OLRFs. One possible mechanism is to extract precise object location information from the overlapping region of the visual and tactile OLRFs, producing small, intersection-like bimodal OLRFs (Figure 4A, top). Alternatively, neurons may merge visual and tactile OLRFs into large, union-like receptive fields, which could facilitate object detection (Figure 4A, bottom). We observed neurons resembling both hypothetical mechanisms (or integration modes). To quantify this, we computed a union-intersection score (UI score). The UI score measures how similar a bimodal OLRF is to the union or intersection of the tactile and visual OLRFs of the same neuron (Figure 4B). A high UI score indicates a more union-like integration mode, while a low UI score indicates a more intersection-like mode. Many bimodal OLRFs were similar to neither union nor intersection patterns, and were also poorly matched to the corresponding tactile or visual OLRFs (Figure S3A, S3B). These neurons were excluded from subsequent analyses using a similarity score, as illustrated in Figure 4B (see Methods).

**Figure 4.**
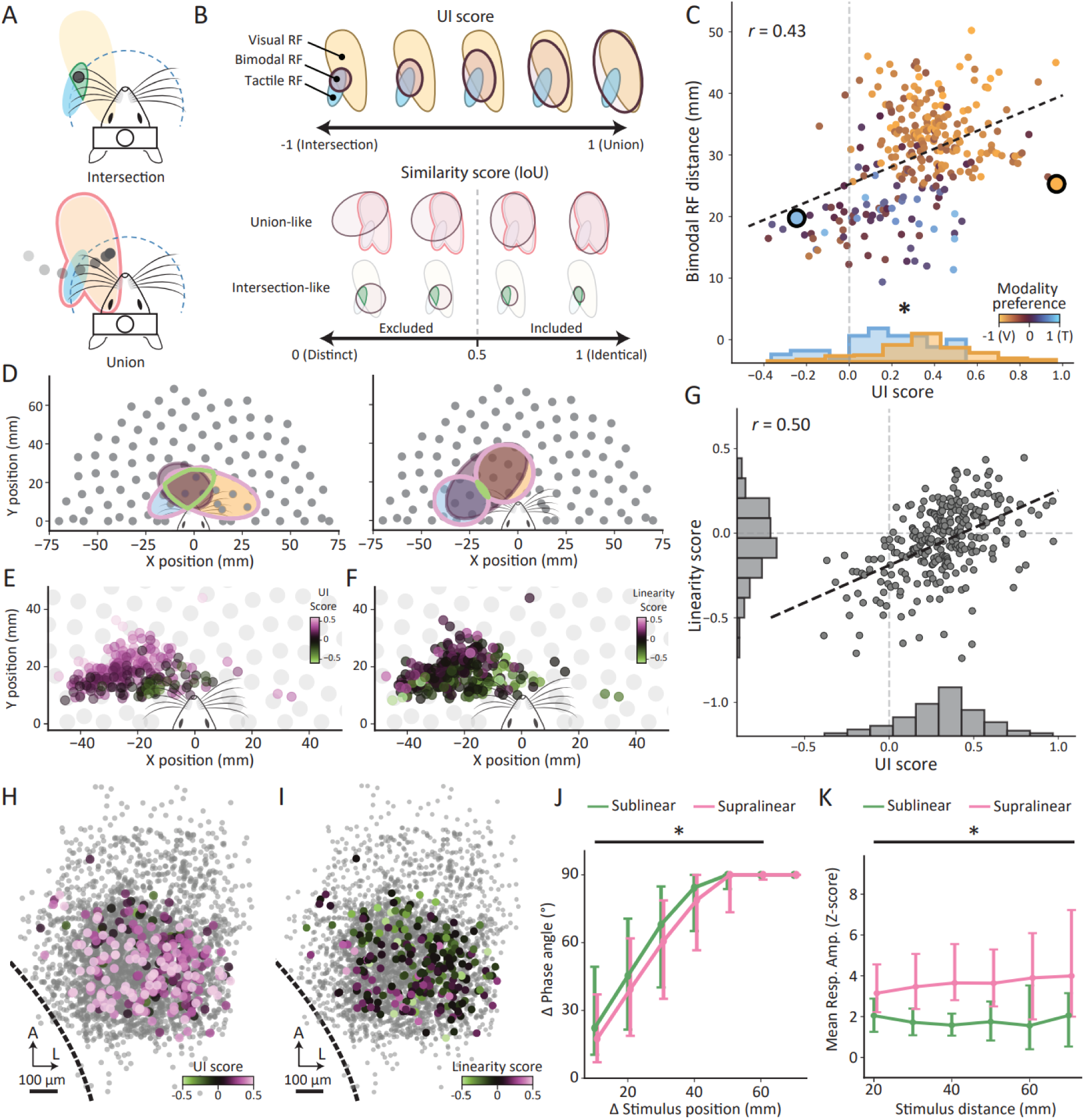
Visuo-tactile integration modes exhibit distance-dependent organization and computational strategies. **A.** Conceptual framework illustrating two multisensory integration modes: *Intersection* (bimodal OLRFs span only regions of overlap between visual and tactile OLRFs) versus *Union* (bimodal OLRFs span combined visual and tactile OLRFs). **B.** Upper: the Union-Intersection score (UI score) quantifies the degree of integration mode preference, ranging from −1 (pure intersection) to +1 (pure union). Lower: the similarity score of bimodal OLRF to each integration mode quantified using the “Intersection over Union” (IoU) metric (see Methods). Only neurons with similarity scores higher than 0.5 were included for the UI score related analysis (227 neurons from 7 mice); **C.** Integration mode preference quantified by UI score increases with preferred object location distance (Pearson r = 0.43, p < 0.001). Dot colors indicate individual neuron modality preferences. Black dashed line: linear regression; gray dashed line: UI-score = 0. Bottom histograms showed a significantly higher UI score of visual dominant neurons (modality preference > 0; yellow) than the score of tactile-dominant neurons (modality preference < 0; blue); *p < 0.001, Mann-Whitney U test. **D.** The bimodal (dark brown), visual (yellow) and tactile (blue) OLRFs of the two example neurons highlighted in **C**. Green and pink outlines demarcate intersection and union areas, respectively. Gray dots show stimulus position grid. **E-F** Scatter plots of bimodal OLRF centers colored by corresponding UI scores (E) or linearity score (F, −1 to 1: sublinear to supralinear) across egocentric near space. **G.** Scatter plots of integration linearity and integration mode preference of bimodal neurons. The regression line (black dash) shows the best fit to the data (Pearson r = 0.50, p < 0.001). Marginal histograms show distributions of linearity scores (left) and UI scores (bottom). Gray dashed lines: zero values of linearity and UI score. **H-I.** Cortical topography of UI score (**H**) and linearity score (**I**) across RL. A, anterior; L, lateral. Black dashed lines indicate RL-V1 boundaries. **J.** Difference in stimulus position plotted against phase angle difference of the corresponding population vectors for sublinear (green) and supralinear (magenta) neurons. Asterisks indicate significantly higher phase angle differences in sublinear compared to supralinear neurons (p < 0.001, Mann-Whitney U test with Bonferroni correction) in the corresponding distance bin of stimulus position separation. **K.** Stimulus distance plotted against the mean response amplitude of the responding sublinear and supralinear neurons (response amplitude > 0). Asterisks indicate significantly higher response amplitudes in supralinear neurons (p < 0.001, Mann-Whitney U test with Bonferroni correction) in the corresponding stimulus distance bin. The nodes and error bars in **J**-**K** correspond to median and the 25^th^ −75^th^ percentiles.

Overall, neurons displayed diverse integration modes, spanning the spectrum from intersection-like to union-like integration (Figure 4C). Crucially, neurons’ integration modes were not randomly distributed across the population. Rather, we found a strong correlation between the integration mode and OLRF distance in mouse near space: neurons with proximal bimodal OLRFs were more intersection-like, while those with distal bimodal OLRFs were more union-like (Figure 4C; example neurons shown in 4D). This relationship produced a spatial gradient of integration mode across contralateral near space, with intersection-like integration predominating near space accessible to both whiskers and vision, and union-like integration in far space (Figure 4E). In addition, bimodal RFs with high similarity scores to union or intersection patterns were more spatially concentrated than the overall population towards a region of the mouse near space, which is well within reach of both vision and whiskers (Figure S3C). These results highlight distance as a critical factor in shaping visuo-tactile integration in area RL.

While the integration mode captures differences in the spatial extent during visuo-tactile integration, integration linearity reflects changes in response magnitude. Supralinear integration means that the response amplitude to the bimodal stimulus is higher than the sum of unimodal responses, while sublinear integration means a lower bimodal response amplitude than the sum. We quantified integration linearity by comparing bimodal response to summed unimodal responses^13^ (Figure 4F). Consistent with previous findings^13,18^, RL neurons displayed diverse levels of integration linearity, spanning from sublinear to supralinear (Figure 4F-G). Notably, the linearity gradient closely resembles the integration mode gradient, showing a shift from sublinear in the near region to supralinear in the farther region in mouse near space (Figure 4F). This similarity was confirmed by a strong correlation between integration mode and linearity (Figure 4G). Both gradients were also seen in the cortical organization of RL: neurons near V1 were more supralinear and union-like, while those farther from V1 were more sublinear and intersection-like (Figure 4H-I). A similar linearity gradient was observed in the spatial distribution of 3D OLRFs, indicating the distance-linearity relationship is not limited to the 2D plane, but extends to three-dimensional space (Figure S3D-E).

To understand how the spatial organization of integration linearity shapes the population code for object location, we split the multimodal neurons into sublinear and supralinear groups. We then computed the population response vector for each group and stimulus location, with each experiment session analyzed separately. A large phase angle between response vectors for different stimulus locations indicates more distinct population representations, independent of overall response amplitude. Using this metric, we found that the phase angle difference was significantly larger in the sublinear group than in the supralinear group, exclusively for bimodal stimuli (Figure 4J & S4A). This result suggests that sublinear neurons provide a more distinguishable code for object location than supralinear neurons when objects are accessible by both vision and whiskers. In contrast, both the average response amplitude and amplitude modulation were significantly lower in the sublinear group than in the supralinear group, again exclusively for bimodal stimuli (Figure 4K & S4B-D). These observations rule out the possibility that the lower phase angle difference in the supralinear group is due to weaker responses. Together, these results suggest that sublinear neurons encode object location more precisely, while supralinear neurons contribute a more salient population code for the overall presence of objects in near space.

## Discussion

In this study, we employed fine-grain multimodal stimulation to map object location receptive fields (OLRFs) across visual and tactile modalities in a subregion of mouse posterior parietal cortex, area RL. Using this approach, we systematically assessed how neurons in this area encode and integrate object location in egocentric reference frames during active sensing. By comparing visual, tactile, and bimodal OLRFs, we observed distance-dependent visuo-tactile integration across neurons and OLRF properties. Such dependence appears as a cortical gradient in RL, with which visual and tactile topographic maps partially overlap. This distance-dependency profoundly shapes visuo-tactile integration as both integration mode (how inputs are spatially combined) and linearity correlate with distance. This results in distinct sublinear and supralinear neural codes for object location and object detection.

Previous studies reported partially co-aligned visual and tactile topographic maps in RL^13,16,17^. This co-alignment could have indicated either a shared egocentric reference frame across modalities or a similar representation of upper and lower space along retinotopic and somatotopic maps. In this study, we show that most OLRFs are additionally tuned to the dimension absent in the two-dimensional retinotopic/somatotopic framework, namely distance (Figure 2). This distance tuning is prevalent across tactile, visual, and bimodal OLRFs, supporting the concept of a shared egocentric reference frame for encoding mouse near-space. This idea is supported by findings made in PPC regions with similar function and location in other mammalian species: In rodents, a significant proportion of PPC neurons encode egocentric object locations using vision alone^2,21^. In primates, area VIP (ventral intraparietal) forms multisensory egocentric representations of peri-head space^6,7,12,22,23^.

While visual and tactile topographical maps are partially aligned in RL, they diverge in regions dominated by the other modality (Figure 3). These unimodal-dominated regions, close to V1 and the barrel cortex, respectively, sample near-space regions that reflect the accessible range of each modality, vision for farther and whiskers for closer space. This indicates that visuo-tactile integration in RL is not simply resulting from merged retinotopic and somatotopic maps. Rather, it suggests a complex organization connecting modality accessibility to spatial distance and engaging multiple cross-modal circuits, as suggested by recent studies in which tactile stimulation may drive or suppress visual cortex activity^14,24^.

This distance influence extends beyond receptive field properties to the mode and linearity of visuo-tactile integration (Figure 4). At close distances, in regions accessible to both visual and tactile inputs, neurons tend to integrate inputs from overlapping regions sublinearly; at farther distances, where overlap is reduced, neurons often integrate input from the combined regions supralinearly. This finding is in line with the broadly observed principle of inverse effectiveness across neural correlates, species and modalities^13,25–28^. This principle states that multisensory enhancement is more pronounced when the unimodal inputs are weakly activated^27^. We further found that the spatial extent of the integration covaries with the degree of multimodal enhancement. During visuo-tactile integration, sublinear neurons more precisely encode object location, while supralinear neurons respond strongly to the mere presence of an object. These distinct integration modes suggest a possible distance-dependent shift in the ethologically relevant computational goals: precise object manipulation (e.g. prey capture) is important at very close distances, while object detection is more relevant at farther distances.

## Acknowledgement

We thank M. Sperling, M. Linnenbrink and F. Voss for technical assistance; P. Goltstein for help at various stages of the project; J. Hinz for discussions. This project was funded by the Max Planck Society.

## Author contributions

Y.Z., and M.H. designed experiments. Y.Z. and U.L. conducted experiments. Y.Z. built the stimulus setup and analysis pipelines. Y.Z., U.L., A.L., T.B. and M.H. discussed the data and wrote the manuscript.

## Declaration of AI-assistance in the writing process

The authors declare that any use of AI tools was limited to grammar checking and readability improvements. The authors reviewed, edited, and take full responsibility for the content of the published article.

## Methods

### Lead Contact

Further information and requests for resources and reagents should be directed to and will be fulfilled by the lead contacts, Yue Zhang and Mark Hübener.

### Experiment Model and Subject Details

#### Mice

All animal handling and experimental procedures followed the ethical guidelines established by the Max Planck Society and protocols approved by the Regierung von Oberbayern (Beratende Ethikkommission nach §15 Tierschutzgesetz). A total of seven 9–25-week-old female wild-type C57BL/6J mice were used. Mice were housed at 55 ± 5 % humidity and 22 ± 1.5 °C under a 12-hour inversed light-dark cycle with unrestricted access to food and water. Housing consisted of large enclosures (GR900, Tecniplast) enriched with running wheels, tunnels, and shelters. Animals were housed in groups, with temporary individual housing only during post-surgical recovery periods.

No statistical methods were used to predetermine sample size. Since all animals belonged to one experimental group, randomization was not applicable and investigators were not blinded to experimental conditions during data collection and analysis.

### Method details

#### Cranial window implantation

To measure neural activity in area RL using two-photon calcium imaging, cranial window implantation was performed on 5–8-week-old female mice anesthetized via intraperitoneal injection of a mixture of Fentanyl (0.05 mg/kg), Midazolam (5 mg/kg), and Medetomidine (0.5 mg/kg). Anesthetic depth was regularly monitored using the toe pinch reflex. When reflexes were observed, 25% of the original dose of the anesthetic solution was administered subcutaneously to maintain anesthesia. All surgical instruments were heat-sterilized and cleaned with ethanol. Body temperature was maintained with a heating pad, and the eyes were protected with eye cream (Isopto-Max).

Before the initial scalp incision, mice received a general analgesic (Carprofen, 0.5 mg/kg, subcutaneous injection). After disinfection with iodine and ethanol and application of Lidocaine as a local anesthetic, the scalp together with the periosteum was removed. A custom aluminum head bar was fixed to the skull using superglue (Pattex Ultra Gel) and dental cement (Paladur). To improve adhesion, the skull was roughened with a scalpel. Dental cement was also applied around the edge of the scalp wound and over the skull, except for the area of the planned craniotomy.

#### Intrinsic optical imaging guided virus injection

To enhance the accuracy of cranial window placement and virus injections, intrinsic optical imaging was performed through the skull to locate V1. The imaging system consisted of tandem 135 mm f/2.0 and 50 mm f/1.2 lenses, a removable 700-740 nm bandpass filter, and a pco.edge 4.2 LT sCMOS camera. The brain was illuminated with 530 nm light to acquire images of the cortical blood vessel pattern, then with 735 nm light for functional imaging, with the focal plane adjusted to ∼250-500 μm below the cortical surface. Drifting gratings (2 Hz, 0.04 cyc/°; 8 directions changing every 0.6 s, each direction for 7 seconds) covering 0–30° azimuth of the contralateral visual field were presented on a screen 16 cm in front of the mouse 7-10 times. Eye cream was removed from the contralateral eye during stimulation. V1 was located using intrinsic signals in response to visual stimuli, analyzed with a custom MATLAB script.

A circular craniotomy (4 mm diameter) centered on the binocular region of V1 in the right hemisphere was made using a dental drill. The dura mater was carefully removed to expose the cortex. Four to six sites within the binocular region of V1 and the neighboring rostral-lateral area (RL) were selected for virus injections. At each site, 150–200 nL of virus (AAV2/1.Syn.mRuby2.GSG.P2A.GCaMP6s.WPRE.SV40, 1.7 x 10^13^ genome copies/mL) were injected using a glass pipette (tip diameter: 25–35 μm) at a depth of 300–400 μm below the cortical surface. Injections were performed with a pressure micro-injection system (Toohey Company), controlled by a pulse generator (Master-8; 30–40 psi, 20–40 ms pressure pulses at 0.8 Hz). The pipette remained in place for ∼5–10 min after each injection to allow sufficient viral diffusion.

After completing all injections, the craniotomy was closed with a 4 mm glass coverslip, with its rim sealed with superglue. Dental cement was applied to the remaining exposed skull to secure the coverslip. Mice then received subcutaneous injections of 330 μl 5% glucose saline and anesthetic antagonists (Naloxone (1.2 mg/kg), Flumazenil (0.5 mg/kg), and Atipamezole (2.5 mg/kg)). Animals were placed in warm recovery environments in their home cages, and maintained under a reversed 14/10 h light-dark cycle to facilitate imaging during their active (dark) phase.

#### Retinotopic mapping with intrinsic optical imaging

Intrinsic optical imaging was repeated at least 2 weeks after virus injections to visualize cortical retinotopic maps using a drifting flickering checkerboard stimulus as previously described^29^. The imaging procedure followed the same protocol as described in the previous section, except animals were only lightly anesthetized via intraperitoneal injection (Fentanyl (0.035 mg/kg), Midazolam (3.5 mg/kg), and Medetomidine (0.35 mg/kg)).

#### In vivo two-photon calcium imaging

Two-photon imaging was performed on awake animals 4–17 weeks after cranial window implantation. Animals were head-fixed on top of an air-suspended Styrofoam ball (20 cm diameter), allowing them to run freely during imaging. A custom-built two-photon microscope was used for image acquisition^30^. The microscope consisted of a Ti:Sapphire laser with a DeepSee pre-chirp unit (Spectra Physics MaiTai eHP), an 8-kHz resonant galvanometer scanner, and a Nikon 16x objective (0.8 NA). The objective was immersed in ultrasonic gel applied over the cranial window implant. Imaging was performed at a depth of 150–350 μm below the cortical surface, corresponding to layer 2/3 of the cortex. An excitation beam at 940 nm with 10–30 mW laser power (measured under the objective) was used to scan a field of view of 375 x 605 μm at 14.4 Hz with 0.5 x 0.55 μm/pixel resolution^2^. Custom LabVIEW and Python software were used for image acquisition and synchronization with the stimulus system.

#### Motorized multimodal stimulus system

A custom motorized stimulus system was developed for precise presentation of visual and/or tactile stimuli at defined spatial locations within the peri-facial near space of head-fixed mice during two-photon calcium imaging. This system enabled systematic mapping of object location receptive fields (OLRFs) across different sensory modalities. The system consisted of a three-axis motorized stage, a stimulus head, and a motorized whisker mask.

The motorized stage used programmable servo motors to drive movement along X, Y, and Z axes (X and Y axes: Dynamixel XC330; Z axis: Dynamixel XL330, Robotis), with force transmitted through a custom 3D-printed gear assembly. Each axis was constrained by ball-bearing rail guides to ensure smooth, linear motion. Motor control was achieved via the Dynamixel U2D2 USB interface, enabling high-speed serial communication with a host computer running custom Python software based on the official Dynamixel SDK for real-time position control and feedback. Positioning accuracy was 0.5 mm for all axes.

The stimulus head was equipped with white LED-coupled optical fibers and a physical object, allowing synchronized presentation of visual and tactile cues. For 2D mapping, the stimulus consisted of a vertical bar (40 mm length x 2.2 mm diameter) illuminated by white LED light. For 3D mapping, a small sphere (6 mm diameter) replaced the vertical bar stimulus. Independent presentation of visual and tactile cues was achieved as follows: (1) for visual-only stimulation a fine-net whisker mask covered the mouse’s face to prevent tactile contact while allowing visual stimulation; (2) for tactile-only stimulation the LED illumination was turned off, preventing visual stimulation while permitting whisker contact; (3) for bimodal stimulation both visual and tactile cues were present. The whisker mask was attached to a carbon fiber rod, controlled by a servo motor, rotating to cover the mouse face during deployment and retracting out of the stimulation area when not needed, with ∼3-second deployment and retraction times.

Spatial coordinates of the stimulus were defined with X, Y, and Z axes corresponding to left-right, front-back, and up-down directions, respectively. The coordinate system origin was set at the midpoint between the eyes and nose, individually calibrated for each mouse before experimentation.

#### Continuous stimulus protocol

A continuous stimulus protocol was designed to enable efficient computation of object location receptive fields (OLRFs) across all modality combinations at high spatial resolution.

For 2D mapping, stimulus locations were distributed using Fibonacci sampling across a hemi-circular area (70 mm diameter) centered on the mouse, resulting in 100 positions with 8.88 ± 1.71 mm spacing between neighboring locations. Positions closer than 15 mm to the mouse were excluded as a collision protection zone. For 3D mapping, 288 stimulus location with 11.03 ± 1.28 mm spacing were arranged in a grid pattern throughout the volume within the distance range 16.91 to 69.86 mm. Due to ongoing optimization of setup precision, stimuli were presented at slightly different locations in some animals, resulting in average shifts of 0.19 mm (2D) and 5.43 mm (3D) from target stimulus locations.

Experiments were conducted with a fixed sequence of conditions for both 2D and 3D stimulation protocols: bimodal, tactile-only, and visual-only stimulation, with 10-second intervals between conditions. Each condition was repeated for five (2D stimuli) or three trials (3D stimuli), with each trial using a unique trajectory through the same set of spatial locations. Trajectories were generated using a heuristic traveling salesman algorithm to minimize spatiotemporal correlations. During each trial, the stimulus cue moved continuously between locations following the assigned trajectory, with no settling time at any position. The travel time between consecutive positions was 0.5 seconds for 2D stimuli and 0.4 seconds for 3D stimuli. Synchronization between stimulus position and imaging was maintained with 2 ms accuracy. The protocol completed 2D mapping in less than 14 minutes and 3D mapping in less than 18 minutes.

### Quantification and statistical analysis

#### Calcium imaging data preprocessing

All data preprocessing and analysis were performed in Python. Calcium movies were processed using Suite2p for motion correction^31^. Neurons were detected from maximum projection images and selected as regions-of-interest (ROI) for calcium trace extraction by Cellpose^32^. Calcium traces were extracted by summing pixel fluorescence intensities within each ROI across all frames and normalized by z-scoring:

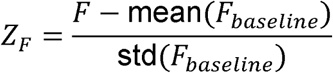

where *F* is the raw fluorescence trace, and *F_baseline_* is the fluorescence signal during a ∼20-second baseline period recorded once before stimulation started. Normalized calcium signals were synchronized with stimulus position data for subsequent analyses.

To combine data from multiple imaging sessions within the same animal, we manually aligned average calcium images using blood vessel landmarks. Displacement vectors were computed between each session and a reference session, then applied to shift ROI coordinates onto a unified spatial map. Potentially duplicated ROIs from overlapping fields of view were not excluded, as arbitrary exclusion criteria could introduce a greater bias than the inclusion of duplicates.

#### Reverse-correlation analysis

OLRFs were computed by reverse-correlating z-scored calcium activity traces with stimulus trajectories across all trials for each condition as follows:

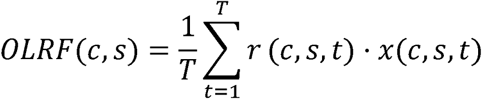

where c indicates the stimulus condition (visual-only, tactile-only, or bimodal), s is the spatial location, and T is the total number of time points for each condition and stimulus location. *x*(*c*, *s*, *t*) indicates stimulus presence at location s at time t under the condition c, and *r*(*c*, *s*, *t*) is the corresponding calcium response at that time point.

To combine data across animals, actual stimulus positions were interpolated to match standard protocol positions using linear interpolation for animals with slightly deviating stimulus protocols. The maximum fractional loss in signal correlation 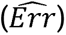 due to spatial interpolation was estimated as:

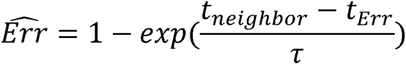

This formula compares the estimated travel time for spatial interpolation deviation (*t_Err_*: 0.011 s for 2D, 0.19 s for 3D) with travel time for neighboring stimulus locations (*t_neighbor_*: 0.5 s for 2D, 0.4 s for 3D), scaled by the calcium signal decay time constant (τ: 2.59 s)^33^. Since calcium signal correlations decay exponentially over time, larger spatial deviations (longer *t_Err_*) lead to greater fractional loss in signal correlation. The maximum fractional losses in signal correlation were 0.43% for 2D and 7.05% for 3D, indicating that spatial interpolation introduces negligible error for OLRF estimation and subsequent analyses.

Given the strong autocorrelation in the stimulus positions and the slow dynamics of calcium signals, temporal delays between stimulus presentation and neural responses were considered negligible, and no delay correction was applied.

#### Significant OLRF detection and statistical analysis

Significant object location receptive fields were identified using a two-step statistical framework: first determining whether neurons exhibited spatial structure, then identifying specific regions of significant response clustering.

##### Step 1: Spatial Autocorrelation Testing (Moran’s I)

Each neuron was tested for receptive field structure using Moran’s I statistic^34^. This test determined whether the spatial response pattern showed significant spatial autocorrelation, as receptive field structures are assumed to be locally coherent rather than scattered and fractal. The Moran’s I statistic was calculated as:

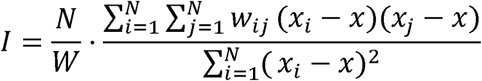

where *N* is the number of spatial locations, *x_i_* is the Ca^2+^ response at location *i*, *x* is the mean response across all locations, and *w_ij_* represents the spatial weights matrix. Spatial neighborhood structure was defined using distance-based Gaussian weights with σ = 10 mm.

Statistical significance was assessed through permutation testing with 1000 random permutations of the neural response vector, generating an empirical null distribution for each neuron. Neurons with significant spatial autocorrelation (p < 0.05 with Benjamini-Hochberg FDR correction across neurons) were classified as having detectable receptive field structures.

##### Step 2: Local Spatial Clustering (Getis-Ord G_i_*)

For neurons passing the global autocorrelation test, specific spatial regions of significantly elevated neural responses were identified using the local Getis-Ord G_i_* statistic^35^. This analysis revealed spatial clusters of strong responses corresponding to significant OLRF components. The G_i_* statistic was computed for each spatial location *i* as:

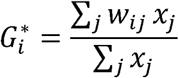

The same distance-based Gaussian weight matrix (σ = 10 mm) defined local neighborhoods. Statistical significance of local clusters was determined using the same permutation testing framework (1000 permutations) with a p < 0.05 threshold. The union of all spatial locations with significant local clustering (G_i_*p < 0.05) defined the boundaries of significant OLRFs. Only neurons with significant spatial autocorrelation containing at least one significant local cluster were included in subsequent analyses.

#### OLRF fitting and analysis

To quantify OLRF geometry and orientation, ellipses (2D) or ellipsoids (3D) were fitted to significant OLRFs using weighted principal component analysis (wPCA)^36^. This approach weights each spatial location by its mean Ca^2+^ response to account for varying response intensities across OLRF locations.

The fitting procedure involved constructing a weighted covariance matrix from the spatial coordinates of all significant locations within each OLRF. Weights corresponded to mean Ca^2+^ responses at those locations averaged across all trials. The weighted covariance matrix was computed as:

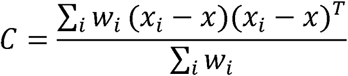

where *w_i_* is the mean Ca^2+^ response weight at location *i*, *x_i_* is the 2D or 3D coordinate vector, and *x* is the weighted centroid. Principal components were extracted via eigen-decomposition, with eigenvectors scaled to cover 90% of the weighted data distribution. The resulting ellipses or ellipsoids were centered at the weighted centroid and oriented along the principal component axes.

Receptive field orientation was defined by the phase angle of the major axis for 2D OLRFs. OLRF size was quantified using ellipse area (*A*) or ellipsoid volume (*V*):

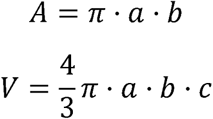

where a, b, and c are the lengths of the semi-major and semi-minor axes.

#### OLRF feature map computation and alignment

OLRF feature maps were aligned across sessions and animals using a two-step approach. First, OLRF feature maps were manually aligned within each animal across multiple imaging sessions using blood vessel patterns visible in average two-photon images as landmarks as mentioned above. For alignment across animals, azimuth maps computed from OLRF centers served as a common reference frame. The coordinate of the median location of neurons tuned to the medial visual field (indicated by OLRF center azimuth > −15 degrees) was computed. As confirmed by comparison with retinotopic maps from intrinsic optical imaging, these neurons were distributed near the boundary between area RL and V1. All OLRF feature maps were then translated to align these reference points across animals.

To visualize the spatial distribution of OLRF features, smoothed feature maps were computed using a Gaussian kernel (σ = 50 μm) to convolve ROIs on a merged feature map labeled with the corresponding OLRF property value. To reduce the influence of non-uniform ROI distributions, regions with fewer than 10 ROIs within a 90 μm radius were excluded.

For each bimodal feature map, a principal gradient axis was determined by computing the most dominant descending direction of the gradient from the feature map smoothed with a larger Gaussian kernel (σ = 100 μm, indicated by the black arrows in Figure 3E-H). This principal axis was used to compute averaged feature values along the axis for the analyses used in Figure 3I-L.

#### Modality preference and integration linearity score

The modality preference score was computed for each neuron as described^16^:

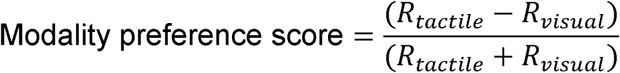

where *R_tactile_* and *R_visual_* are the mean Ca^2+^ responses within the tactile and visual OLRFs, respectively. For neurons with significant OLRFs in only one modality, extreme index values were assigned: +1 for neurons with only tactile OLRFs (tactile-only responsive) and −1 for neurons with only visual OLRFs (visual-only responsive).

The integration linearity score was computed as described^13^:

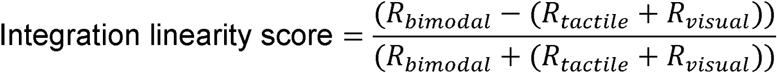

where *R_bimodal_* is the mean Ca^2+^ response within the bimodal OLRF.

#### Union-Intersection (UI) score

To quantify the degree of overlap between visual and tactile OLRFs for individual neurons, we developed a Union-Intersection (UI) score metric. Neurons with significant OLRFs under all three conditions were included for analysis. The union and intersection of the visual and tactile OLRFs were computed for each neuron. We computed a similarity score between bimodal OLRFs (*B*) and the intersection (*I*) or union (*U*) regions using the “Intersection over Union” (IoU) metric as follows:

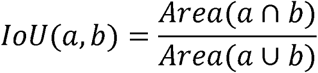

where *a* and *b* are two spatial regions, and *Area*(*a* ∩ *b*) and *Area*(*a* ∪ *b*) are the areas of the intersection and the union of the two regions, respectively. The similarity score of bimodal OLRFs to tactile and visual OLRFs are computed in the same way.

The UI score was then calculated as:

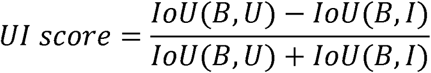

Visual-tactile (VT) scores, evaluating bimodal OLRF similarity to corresponding visual versus tactile OLRFs, were computed in a similar way:

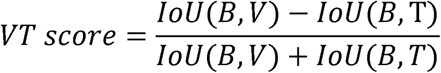

#### Population vector analysis for sublinear and supralinear neurons

To investigate the contributions of sublinear and supralinear neurons to the overall population response, we conducted a population vector analysis. Neurons were categorized as sublinear or supralinear based on their integration linearity scores: neurons with scores less than −0.1 are categorized as sublinear, while the ones with a score larger than > 0.1 are supralinear.

For a stimulus position at location *i*, the population response vector, **P***_i_*, was constructed by concatenating the mean *Ca*^2+^ responses of all sublinear or supralinear neurons at that location:

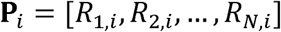

where *R_n,i_* represents the mean *Ca*^2+^ response of neuron *n* at location *i*, and *N* denotes the total number of neurons in the sublinear or supralinear group.

The phase angle difference, Δ*θ_i,j_*, between population response vectors at two stimulus locations *i* and *j* was calculated using cosine similarity:

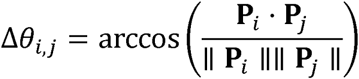

The location information could also be encoded in population responses through gain modulation. To verify this, we computed the mean response amplitudes of the sublinear and supralinear neuron populations for each location. To address potential ambiguities arising from response sparsity and amplitude, these mean response amplitudes were calculated using only responsive neurons with mean activity z-scores greater than 0 at the respective location.

## Supplementary Figures

**Figure S1.**
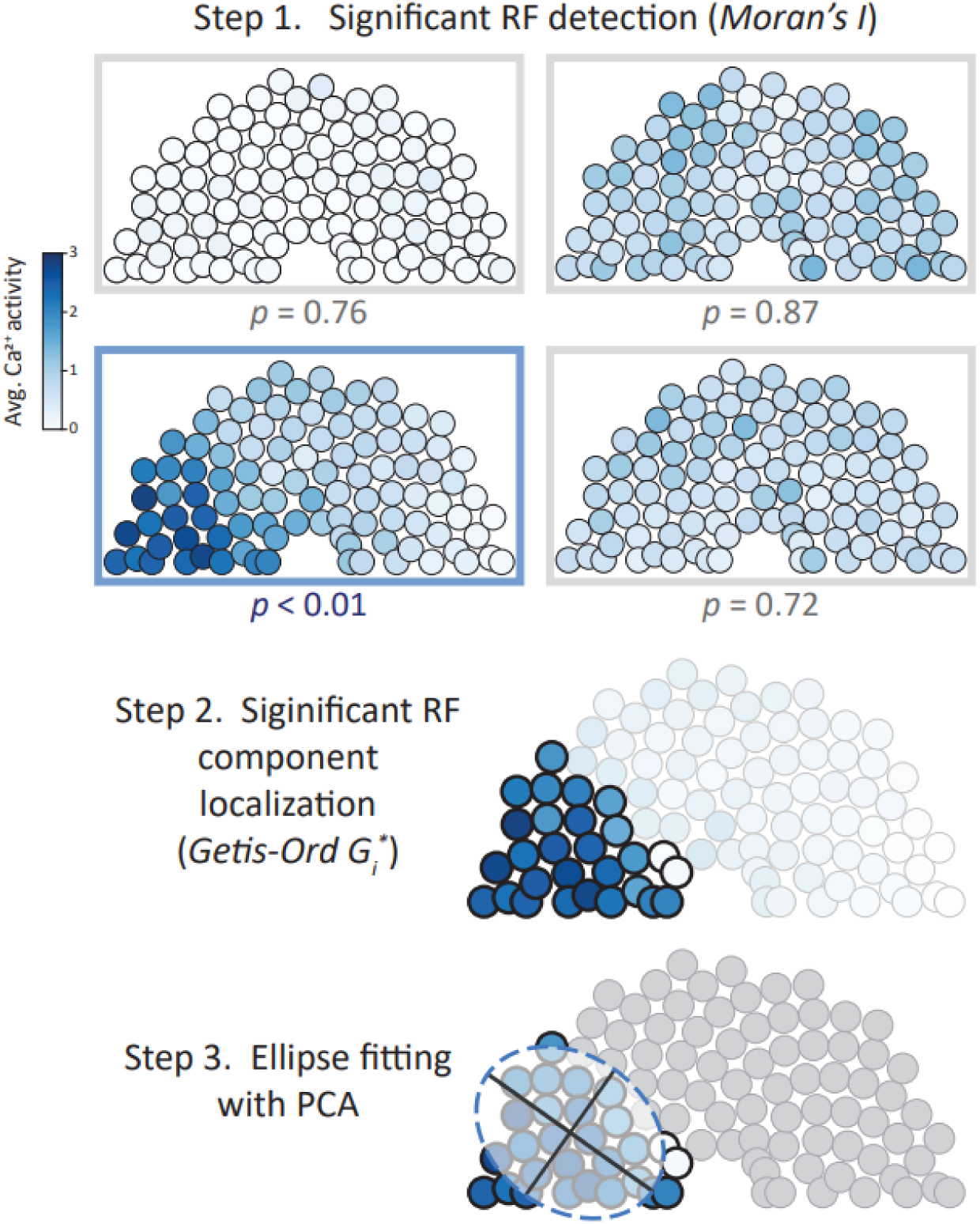
Two-step statistical framework for significant OLRF detection. **Step 1.** One of the four example OLRFs exhibited spatial structure (blue box), which was detected with Moran’s I test. **Step 2.** The significant receptive field (RF) component within this OLRF was localized with Getis-Ord G_i_* in the second step. **Step 3.** An ellipse was fitted to the significant OLRF using weighted PCA. See Methods for details.

**Figure S2.**
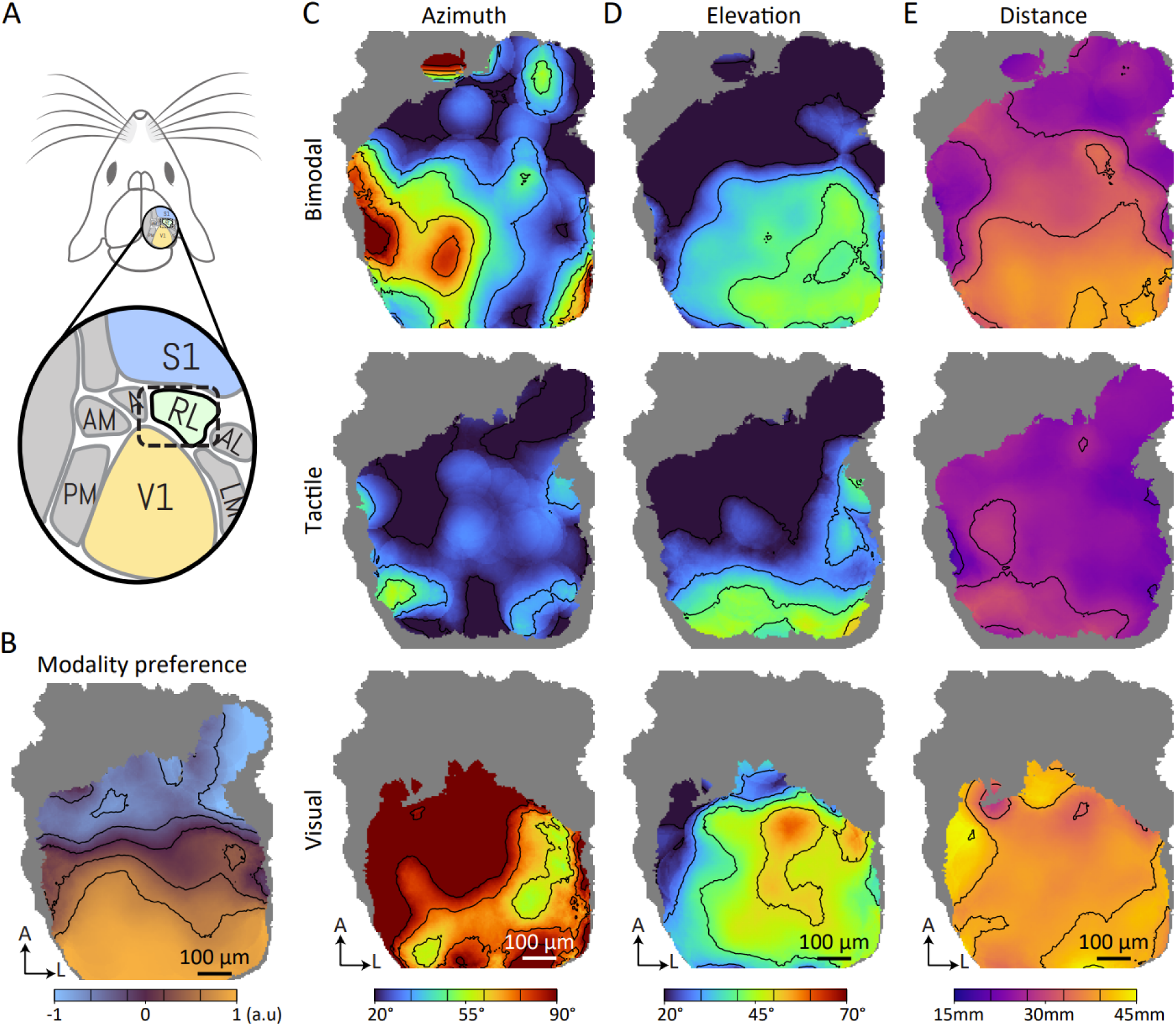
Smoothed feature maps of 3D OLRFs. **A.** Schematic of the recorded area in mouse visual cortex. **B.** Smoothed modality preference maps across RL. Gray regions indicate insufficient data (n < 10 cells within the 180 μm neighboring region). Contour lines indicate iso-modality-preference-score boundaries at intervals marked on the color bar. **C-E.** Smoothed feature maps of OLRF azimuth (**C**), elevation (**D**) distance (**E**) under bimodal (top), tactile (middle) and visual (bottom) stimulus conditions across RL. Contour lines indicate iso-value boundaries at intervals marked on the corresponding color bars.

**Figure S3.**
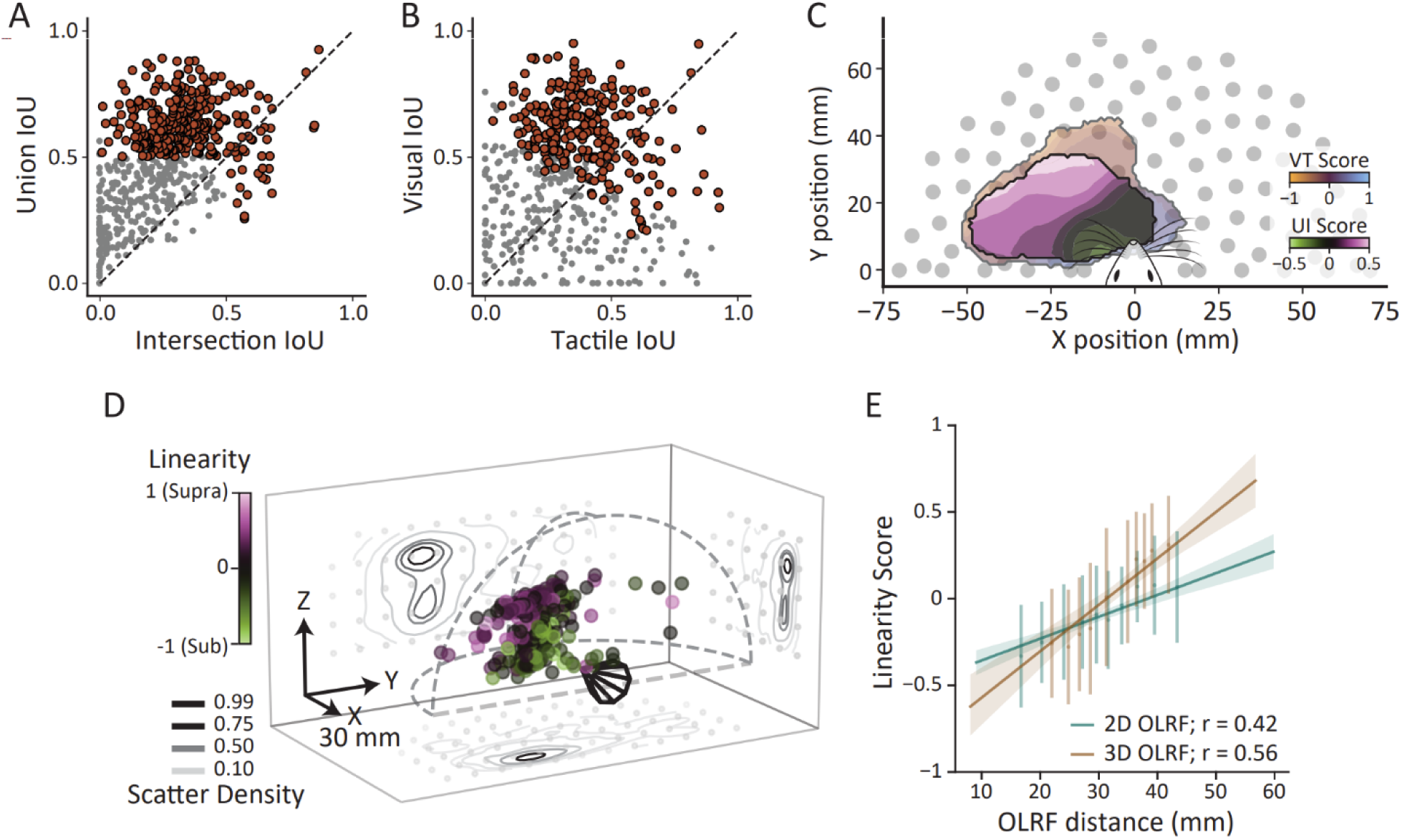
Integration mode and linearity analysis of 2D and 3D OLRFs. **A.** Bimodal OLRF similarity to the intersection of visual and tactile OLRFs (Intersection IoU) versus similarity to their union (Union IoU). **B.** Bimodal OLRF similarity to tactile OLRFs (Tactile IoU) versus similarity to visual OLRFs (Visual IoU). Red symbols in **A-B**: neurons included in UI score calculations (intersection/union IoU > 0.5). **C.** Smoothed spatial distributions of UI and VT scores. Filled contours (green-purple) show local average UI scores computed with Gaussian kernels for neurons highlighted in red (**A-B**). Semi-transparent background contours (brown-blue) indicate local average VT scores for all neurons. **D.** 3D bimodal OLRF centers colored by linearity scores. Cone-shaped polygon indicates mouse head position and orientation. Contour lines represent scatter density estimated with Gaussian kernels (normalized to [0,1], 1 indicates peak density), projected onto cardinal planes (gray dots: stimulus position projections). Dashed arcs outline stimulus field boundaries. **E.** OLRF center distances versus linearity scores for 2D (green) and 3D (brown) data. Solid lines show linear fits with confidence intervals (shaded areas). Points and error bars represent mean linearity score ± standard deviations in each distance bin. Pearson correlations (*r*) are indicated.

**Figure S4.**
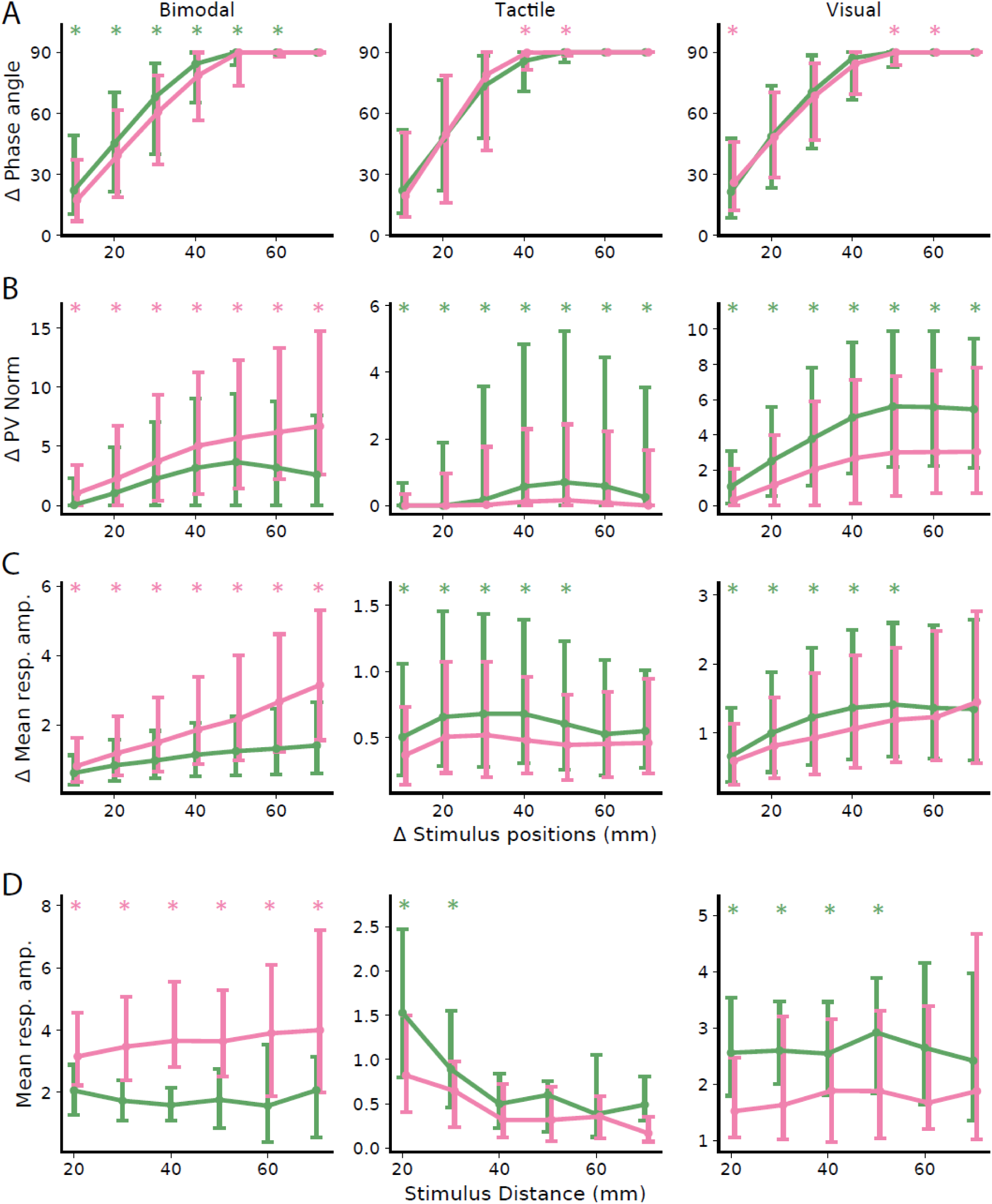
Comparison of population response properties between sublinear and supralinear neurons across stimulus conditions. **A-B.** Changes in phase angles (**A**) and vector norms (**B**) of the population vectors are analyzed as the spatial separation between stimuli increases, comparing how sublinear (green) and supralinear (magenta) neurons encode stimulus positions under bimodal, tactile, and visual conditions. **C.** Differences in the mean response amplitudes of sublinear (green) and supralinear (magenta) neurons (for neurons with amplitude > 0) are compared across increasing spatial separations between stimuli. **D.** Mean response amplitudes of sublinear and supralinear neurons plotted against the distance of stimuli cues across conditions. The nodes and error bars indicate the median and the 25^th^-75^th^ percentiles. Pink asterisks denote bins where the supralinear neuron distribution is significantly higher than the sublinear distribution; green asterisks indicate the reverse (p < 0.001, Mann-Whitney U test with Bonferroni correction).

## References

1. Anzai, A., Ohzawa, I., and Freeman, R.D. (1999). Neural mechanisms for encoding binocular disparity: receptive field position versus phase. J. Neurophysiol. 82, 874–890.

2. La Chioma, A., Bonhoeffer, T., and Hübener, M. (2020). Disparity Sensitivity and Binocular Integration in Mouse Visual Cortex Areas. J. Neurosci. 40, 8883–8899. 10.1523/jneurosci.1060-20.2020.

3. Boone, H.C., Samonds, J.M., Crouse, E.C., Barr, C., Priebe, N.J., and McGee, A.W. (2021). Natural binocular depth discrimination behavior in mice explained by visual cortical activity. Curr. Biol. 31, 2191–2198.e3. 10.1016/j.cub.2021.02.031.

4. Andersen, R.A., Essick, G.K., and Siegel, R.M. (1985). Encoding of Spatial Location by Posterior Parietal Neurons. Science 230, 456–458. 10.1126/science.4048942.

5. Bouvier, G., Senzai, Y., and Scanziani, M. (2020). Head Movements Control the Activity of Primary Visual Cortex in a Luminance-Dependent Manner. Neuron 108, 500–511.e5. 10.1016/j.neuron.2020.07.004.

6. Colby, C.L., Duhamel, J.R., and Goldberg, M.E. (1993). Ventral intraparietal area of the macaque: anatomic location and visual response properties. J. Neurophysiol. 69, 902–914. 10.1152/jn.1993.69.3.902.

7. Colby, C.L., and Duhamel, J.-R. (1996). Spatial representations for action in parietal cortex. Cogn. Brain Res. 5, 105–115. 10.1016/S0926-6410(96)00046-8.

8. Estebanez, L., Bertherat, J., Shulz, D.E., Bourdieu, L., and Léger, J.-F. (2016). A radial map of multi-whisker correlation selectivity in the rat barrel cortex. Nat. Commun. 7, 13528. 10.1038/ncomms13528.

9. Clancy, K.B., Schnepel, P., Rao, A.T., and Feldman, D.E. (2015). Structure of a Single Whisker Representation in Layer 2 of Mouse Somatosensory Cortex. J. Neurosci. 35, 3946–3958. 10.1523/JNEUROSCI.3887-14.2015.

10. Pluta, S.R., Lyall, E.H., Telian, G.I., Ryapolova-Webb, E., and Adesnik, H. (2017). Surround Integration Organizes a Spatial Map during Active Sensation. Neuron 94, 1220–1233.e5. 10.1016/j.neuron.2017.04.026.

11. Mohan, H., De Haan, R., Broersen, R., Pieneman, A.W., Helmchen, F., Staiger, J.F., Mansvelder, H.D., and De Kock, C.P.J. (2019). Functional Architecture and Encoding of Tactile Sensorimotor Behavior in Rat Posterior Parietal Cortex. J. Neurosci. 39, 7332–7343. 10.1523/jneurosci.0693-19.2019.

12. Sereno, M.I., and Huang, R.-S. (2014). Multisensory maps in parietal cortex. Curr. Opin. Neurobiol. 24, 39–46. 10.1016/j.conb.2013.08.014.

13. Olcese, U., Iurilli, G., and Medini, P. (2013). Cellular and Synaptic Architecture of Multisensory Integration in the Mouse Neocortex. Neuron 79, 579–593. 10.1016/j.neuron.2013.06.010.

14. Weiler, S., Rahmati, V., Isstas, M., Wutke, J., Stark, A.W., Franke, C., Graf, J., Geis, C., Witte, O.W., Hübener, M., et al. (2024). A primary sensory cortical interareal feedforward inhibitory circuit for tacto-visual integration. Nat. Commun. 15, 3081. 10.1038/s41467-024-47459-2.

15. Wang, Q., and Burkhalter, A. (2007). Area map of mouse visual cortex. J. Comp. Neurol. 502, 339–357. 10.1002/cne.21286.

16. Guyoton, M., Matteucci, G., Foucher, C.G., Getz, M.P., Gjorgjieva, J., and El-Boustani, S. (2025). Cortical circuits for cross-modal generalization. Nat. Commun. 16. 10.1038/s41467-025-59342-9.

17. Matsumoto, H., Murakami, T., and Ohki, K. (2025). Topographic correspondence between retinotopic and whisker somatosensory map in mouse higher visual area and its development. Front. Neural Circuits 19. 10.3389/fncir.2025.1552130.

18. Nikbakht, N., Tafreshiha, A., Zoccolan, D., and Diamond, M.E. (2018). Supralinear and Supramodal Integration of Visual and Tactile Signals in Rats: Psychophysics and Neuronal Mechanisms. Neuron 97, 626–639.e8. 10.1016/j.neuron.2018.01.003.

19. Schuett, S., Bonhoeffer, T., and Hübener, M. (2002). Mapping Retinotopic Structure in Mouse Visual Cortex with Optical Imaging. J. Neurosci. 22, 6549–6559. 10.1523/JNEUROSCI.22-15-06549.2002.

20. Woolsey, T.A., and Van der Loos, H. (1970). The structural organization of layer IV in the somatosensory region (S I) of mouse cerebral cortex: The description of a cortical field composed of discrete cytoarchitectonic units. Brain Res. 17, 205–242. 10.1016/0006-8993(70)90079-X.

21. Alexander, A.S., Tung, J.C., Chapman, G.W., Conner, A.M., Shelley, L.E., Hasselmo, M.E., and Nitz, D.A. (2022). Adaptive integration of self-motion and goals in posterior parietal cortex. Cell Rep. 38, 110504. 10.1016/j.celrep.2022.110504.

22. Avillac, M., Denève, S., Olivier, E., Pouget, A., and Duhamel, J.-R. (2005). Reference frames for representing visual and tactile locations in parietal cortex. Nat. Neurosci. 8, 941–949. 10.1038/nn1480.

23. Schlack, A., Sterbing-D’Angelo, S.J., Hartung, K., Hoffmann, K.-P., and Bremmer, F. (2005). Multisensory Space Representations in the Macaque Ventral Intraparietal Area. J. Neurosci. 25, 4616–4625. 10.1523/JNEUROSCI.0455-05.2005.

24. Caballero Tapia, A., Cheron, G., Ristori, D., Arckens, L., and Ris, L. (2025). Beyond Vision: Response of the Mouse Visual Cortex to Multimodal Stimulation. Eur. J. Neurosci. 62, e70225. 10.1111/ejn.70225.

25. King, A.J., and Palmer, A.R. (1985). Integration of visual and auditory information in bimodal neurones in the guinea-pig superior colliculus. Exp. Brain Res. 60, 492–500. 10.1007/BF00236934.

26. Alex Meredith, M., and Stein, B.E. (1986). Spatial factors determine the activity of multisensory neurons in cat superior colliculus. Brain Res. 365, 350–354. 10.1016/0006-8993(86)91648-3.

27. Avillac, M., Ben Hamed, S., and Duhamel, J.-R. (2007). Multisensory Integration in the Ventral Intraparietal Area of the Macaque Monkey. J. Neurosci. 27, 1922–1932. 10.1523/JNEUROSCI.2646-06.2007.

28. Truszkowski, T.L., Carrillo, O.A., Bleier, J., Ramirez-Vizcarrondo, C.M., Felch, D.L., McQuillan, M., Truszkowski, C.P., Khakhalin, A.S., and Aizenman, C.D. (2017). A cellular mechanism for inverse effectiveness in multisensory integration. eLife 6, e25392. 10.7554/eLife.25392.

29. Kalatsky, V.A., and Stryker, M.P. (2003). New Paradigm for Optical Imaging: Temporally Encoded Maps of Intrinsic Signal. Neuron 38, 529–545. 10.1016/S0896-6273(03)00286-1.

30. Leinweber, M., Zmarz, P., Buchmann, P., Argast, P., Hübener, M., Bonhoeffer, T., and Keller, G.B. (2014). Two-photon Calcium Imaging in Mice Navigating a Virtual Reality Environment. J. Vis. Exp. JoVE, 50885. 10.3791/50885.

31. Pachitariu, M., Stringer, C., Dipoppa, M., Schröder, S., Rossi, L.F., Dalgleish, H., Carandini, M., and Harris, K.D. (2017). Suite2p: beyond 10,000 neurons with standard two-photon microscopy. Preprint at bioRxiv, 10.1101/061507 https://doi.org/10.1101/061507.

32. Stringer, C., Wang, T., Michaelos, M., and Pachitariu, M. (2021). Cellpose: a generalist algorithm for cellular segmentation. Nat. Methods 18, 100–106. 10.1038/s41592-020-01018-x.

33. Chen, T.-W., Wardill, T.J., Sun, Y., Pulver, S.R., Renninger, S.L., Baohan, A., Schreiter, E.R., Kerr, R.A., Orger, M.B., Jayaraman, V., et al. (2013). Ultrasensitive fluorescent proteins for imaging neuronal activity. Nature 499, 295–300. 10.1038/nature12354.

34. Moran, P.A.P. (1950). Notes on Continuous Stochastic Phenomena. Biometrika 37, 17–23. 10.2307/2332142.

35. Getis, A., and Ord, J.K. (1992). The Analysis of Spatial Association by Use of Distance Statistics. Geogr. Anal. 24, 189–206. 10.1111/j.1538-4632.1992.tb00261.x.

36. Delchambre, L. (2015). Weighted principal component analysis: a weighted covariance eigendecomposition approach. Mon. Not. R. Astron. Soc. 446, 3545–3555. 10.1093/mnras/stu2219.

